# Long lasting non-cellular reactions in sterile soils recapitulate most of the intermediates of the Krebs cycle

**DOI:** 10.1101/2025.07.30.667751

**Authors:** Clémentin Bouquet, Mounir Traikia, Fanny Perrière, Benoit Kéraval, Gaël Alvarez, Loïc Louis, Martin Leremboure, Guillaume Voyard, Sébastien Fontaine, Anne-Catherine Lehours

## Abstract

Over the past decade, chemical evidence has emerged that non-enzymatic metabolic pathways, such as the Krebs cycle, may have existed before cellular life on the primitive Earth. However, the question of whether non-cellular reactions analogous to cell respiration metabolism are still “active” in today’s biosphere and whether they contribute to CO_2_ emissions in contemporary ecosystems remains open. In the present study, we investigated the long-term fate (> 6 months) of organic substrates supplied in sterilised soils in which cell life was not detectable. Through a series of analytical studies performed on the water-extractable fraction of soil exometabolites using chromatography, mass spectrometry and isotope labelling, we demonstrate that sterile soil matrix incubated with [^13^C_6_]-glucose and [^13^C_6_]-citrate can spontaneously generate intermediates of the Krebs cycle, alongside by-products such as acetate and formate. These findings support the hypothesis that extracellular metabolisms (EXOMETs) form a network of non-cellular reactions resembling metabolic pathways involved in aerobic respiration and anaerobic fermentation of cells. This research not only provides insights into the chemical continuity between chemistry and biochemistry but also raises questions about the implications of non-cellular pathways for soil ecosystem functioning and carbon fluxes in the contemporary biosphere.

## Main Text

Metabolism, along with replication and information transfer, is one of the most critical features of living systems. Most of the key reactions in the metabolic network are catalysed by protein enzymes, whose complex molecular structures are the product of Darwinian evolution. This has contributed to the view, which is backed by ‘genetics-first’ theories, that the topologies of metabolic pathways might have developed through the evolution of enzymes. According to this line of thought, there should be no environmental chemistry that mimics metabolism **(1)**.

However, chemical evidence has recently emerged for the existence of non-enzymatic metabolic pathways. One of the first was the serendipitous discovery of glycolysis-like reactions catalysed by the metal cation Fe^2+^ **(2)**. Then, a series of non-enzymatic reactions resembling those of the tricarboxylic acid (TCA) cycle, also known as the Krebs cycle, have been repeatedly documented **(3, 4, 5,6)**, including the identification of TCA intermediates in a carbonaceous meteorite **(7)**. These compelling arguments support that the topologies of the metabolic pathways have their roots in non-enzymatic chemistry **(8)**. From an ecological and biogeochemical perspective, these findings also raise the questions of whether non-cellular reactions analogous to metabolism are still ‘active’ in today’s biosphere, and whether they contribute to respiration and, hence, CO_2_ emissions in contemporary ecosystems.

These questions arise from the recurrent observations of substantial CO_2_ emissions that persist in soils where living biomass has been reduced to undetectable levels by exposure to toxics, gamma irradiation, or autoclaving **(9,10,11)**. Various non-cellular processes have been proposed as potential contributors to CO_2_ emissions in sterile soils including single catalytic reactions driven by reactive oxygen species and/or metals such as iron **(12)** or supported by decarboxylases released during cell lysis **(10,13)**. However, these proposals do not explain how [^13^C_6_]-glucose is completely oxidised to ^13^C-CO_2_ in sterile soils **(10,11,14)**. A decade ago, we proposed the hypothesis of extracellular metabolisms (EXOMETs), consisting of multiple coupled redox reactions that reconstitute an equivalent of a cellular respiration process **(10,11)**. Recently, we have shown that in sterile soils, electron fluxes operate in conjunction with a series of time-ordered reactions controlled by the nature of the substrate, which produce and consume exometabolites and lead to the complete oxidation of organic substrates ([^13^C_6_]- glucose and -citrate) with oxygen as the final electron acceptor. We have also shown that the soil matrix can sustain the complete oxidation of organic compounds for years, a period much longer than the half-life of enzymes, revealing the involvement of stable non-enzymatic catalysts in EXOMET **(14)**. However, to what extent EXOMET activities follow pathways close to central cell metabolism, such as the Krebs cycle, is not yet known.

In the present study, we show that in soils devoid of cell life, citrate and glucose spontaneously generate reactions that recapitulate most of the intermediates of the core biological pathways of the Krebs cycle. In addition, we observe that glucose enhances the synthesis of formate and acetate in sterile soils, supporting the hypothesis that many additional cellular metabolism-like reactions remain to be discovered in contemporary soils.

The analyses were carried out on soil sample extracts obtained from the experimental setup of Bouquet et al. **(14)**. Briefly, soils (pH = 6.2) sterilised by gamma irradiation at 45 kGy and non-sterilised soils were incubated in the dark at 20 °C and a water potential of -100 kPa for 163 days. Five different substrate treatments were applied to sterilised soils: no substrate addition (IS for irradiated soil), the addition of [^13^C_6_]-labelled citrate (^13^C-IS), unlabelled citrate (^12^C-IS), [^13^C_6_]-labelled glucose (^13^G-IS) or unlabelled glucose (^12^G-IS). All treatments were replicated four times (n = 4). The water-extractable fraction of exometabolites with a molecular weight < 3 kDa (hereafter referred to as WEM) was extracted from each replicate sample at each sampling time during the incubation period, which involved sampling of the soils at various intervals (t= 0.2, 3, 6, 17, 23, 66, 100, 163 days). (***SI Appendix***).

First,we analysed WEM using ion chromatography-mass spectrometry (IC-MS) with comparisons against authentic standards to achieve absolute quantification of seven Krebs cycle intermediates **(*SI Appendix*, Figure 1A)**. In non-sterilised soil samples, none of the target Krebs cycle intermediates were detected. By contrast, all of the Krebs cycle target intermediates (citrate, cis-aconitate, α-ketoglutarate, succinate, fumarate, malate and oxaloacetate, **Figure 1A**) were detected in the sterilised soils, with the exception of the citrate which was below the detection threshold at all sampling times for the IS, ^12^G-IS and ^13^G-IS samples. Although the α-ketoglutarate was detected in some samples, its abundance was systematically below the limit of quantification (***SI Appendix*)**. The temporal dynamics of the abundances of the remaining five Krebs cycle intermediates were monitored (**Figure 1A**). It should be noted that ^13^C-labelling did not impact these dynamics since the abundance of the targeted metabolites at each sampling time did not differ significantly between the ^13^C-IS and ^12^C-IS treatments, or the ^13^G- IS and ^12^G-IS treatments. Therefore, to improve statistical robustness, we computed concentrations for each sampling date using the pooled data for each substrate treatment (**Figure 1**).

**Figure 1.**
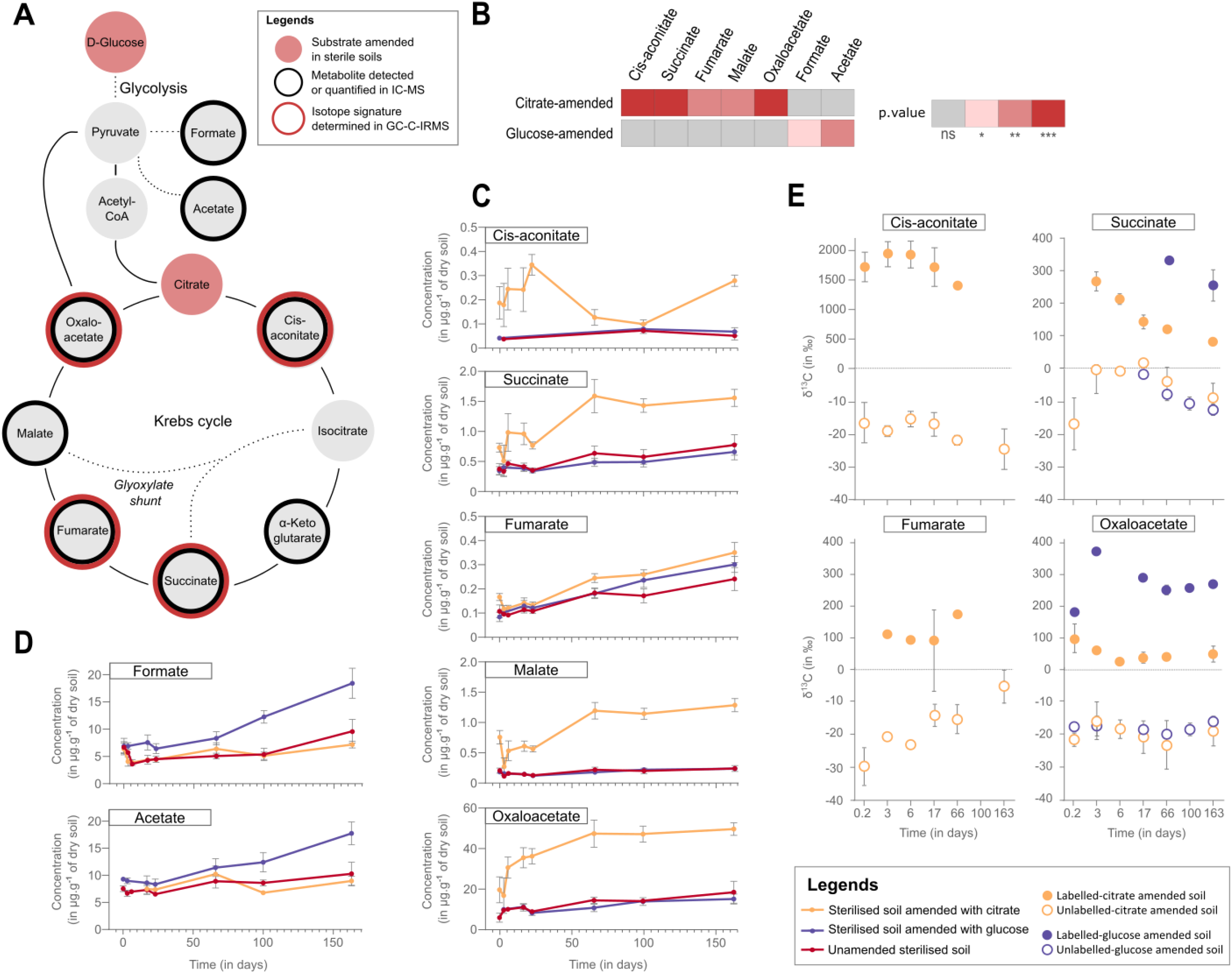
Dynamics of the concentration and carbon isotopic composition of the Krebs cycle intermediates as well as formate and acetate concentrations in sterile soils. **(A)** Schematic representation of the cellular enzyme-catalysed Krebs cycle, as it occurs in yeast, and the glyoxylate shunt (modified from **(8)**), as well as two products (acetate and formate) that can be produced by the fermentation of glucose. The inner (black) and outer (red) circles of the diagram circle indicate the compounds analysed by ionic chromatography coupled to mass spectrometry (IC-MS) and gas chromatography-isotope ratio mass spectrometry (GC-C-IRMS), respectively. Glucose and citrate, coloured pink, were used as substrates in different sterilised soil treatments. **(B)** Quantitative stimulation of Krebs cycle intermediates by citrate or glucose amendment compared to unamended soils. The colour scale represents the results of the two-tailed paired p-value (n=8) for all metabolites, except for cis-aconitic acid for which a one-sample t-test was used (n=8). Asterisks denote p-values as follows: ns: p ≥ 0.05; *: p < 0.05; **: p <0.01; ***: p <0.001. **(C)** Temporal dynamics of the concentrations, analysed by IC-MS, over 163 days of incubation of (from the top to the bottom of the diagram): Cis-aconitate, succinate, fumarate, malate, oxaloacetate in unamended sterilised soils (in red), in glucose-amended sterilised soils (in purple) and in citrate-amended soils (in orange). **(D)** Temporal dynamics of carbon stable isotope ratios (δ^13^C) analysed by GC-C-IRMS of cis-aconitate, succinate, fumarate and oxaloacetate in sterilised soils amended with labelled (solid circles) or unlabelled (open circles) glucose or citrate. For **(C), (D)** and **(E)**, analyses were performed after 0.2, 3, 6, 17, 23, 66, 100, and 163 days of incubation.

During incubation, the production of Krebs metabolites was significantly stimulated (p < 0.01) in sterilised soils that had been amended with citrate, compared to soils that had not been amended and soils that had been amended with glucose. (**Figure 1B**). Although these Krebs compounds exhibited distinct temporal dynamics, they accumulated much more in citrate-amended soils than in the other treatments (**Figure 1C**). For instance, succinate and malate accumulated significantly more (p < 0.01) in citrate- amended soils (1.6 ± 0.1 μg.g^-1^ and 1.3 ± 0.1 μg.g^-1^, respectively) than in unamended soils (0.8 ± 0.2 μg.g^-1^ and 0.2 ± 0.05 μg.g^-1^, respectively) and in glucose-amended soils (0.7 ± 0.1 μg.g^-1^ and 0.2 ± 0.02 μg.g^-1^, respectively) (**Figure 1C**). In contrast, fumarate accumulation was much less pronounced with small differences between treatments (**Figure 1C**). Together with the fact that α-ketoglutarate is below the quantification threshold (***SI Appendix***), these observations suggest a possible shunting of metabolic pathways in sterilised soils with respect to the classical cellular Krebs cycle. Future studies should explore whether non-cellular reactions in sterilised soils also replicate the glyoxylate shunt, a two-step pathway generally regarded as an auxiliary pathway of the Krebs cycle (**Figure 1A**). This is an intriguing prospect, given that Keller et al. **(4)** have demonstrated the ability of non-enzymatic reactions to reproduce the glyxoxylate shunt in experiments simulating the archaean oceans under conditions (e.g. heating to 70°C) that differ greatly from those of the present study **(4, 8)**.

In addition to being the most abundant, oxaloacetate is the metabolite that accumulates the most. By the end of the incubation period, its concentration had reached 49.6 ± 3.1 μg.g^-1^ in citrate-amended sterilised soils (**Figure 1C**). This is more than 2.5 times higher than in sterilised soils or in glucose-amended sterilised soils and between 25 and 130 times higher than the other Krebs metabolites (**Figure 1C**). In the enzyme-catalysed Krebs cycle occurring in cells, oxaloacetate - the end product of a previous round of the cycle - forms citrate by condensing with acetyl-CoA (**Figure 1A**). To date, this step has not been reproduced in non-enzymatic experiments **(4, 8)**. Therefore, the accumulation of oxaloacetate, combined with the lack of citrate detection in unamended and glucose-amended sterilised soils, suggests that oxaloacetate production may represent a dead end for EXOMETs.

As the addition of glucose to sterilised soils did not stimulate the production of Krebs cycle intermediates (**Figure 1B**), we examined the dynamics of acetate and formate, which are considered classic markers of fermentative reactions. The abundances of these two compounds were significantly (p < 0.05) stimulated by the addition of glucose, particularly during the last incubation period (> 66 days, **Figure 1D**). After 163 days, the glucose-amended soils had accumulated 198 % and 173 % more acetate and 256 % and 192 % more formate than the unamended and citrate-amended sterilised soils (**Figure 1D**). These results suggest that EXOMETs may be engaged in additional non-cellular metabolic processes besides aerobic respiration, such as fermentation.

Secondly, to verify that the accumulation of Krebs cycle intermediates was at least partly due to the conversion of labelled or unlabelled glucose and citrate in sterilised soils, the δ^13^C isotope ratios of metabolites in the sterilised soils were measured using gas chromatography-isotope ratio mass spectrometry (GC-C-IRMS) (***SI Appendix***). Each sample was also concurrently analysed using gas chromatography-mass spectrometry (GC-MS) to identify and determine the retention times of the target compounds (***SI Appendix***). Prior to analysis, the samples were derivatised, resulting in the loss of malate during the procedure. In citrate-amended soils, the δ^13^C values of cis-aconitate, succinate, fumarate and oxaloacetate differed significantly (p <0.01) between ^13^C-IS and ^12^C-IS. For cis-aconitate, the average δ^13^C values were 1752.2 ± 283.2 and - 17.5 ± 4.6 in ^13^C-IS and ^12^C-IS, respectively (**Figure 1E**). Due to the low amounts of Krebs intermediates observed in glucose-amended soils, it was not always possible to detect these intermediates by GC-C-IRMS, depending on the metabolite and day of incubation (**Figure 1E**). However, the δ^13^C values of the Krebs intermediates in each measured sample were significantly higher (p <0.001) in ^13^G-IS than in ^12^G-IS showing that glucose also contributes to the formation of succinate and oxaloacetate (**Figure 1E**). This suggests that non-cellular reactions in soils can trigger cascades of reactions capable of replicating all or part of glycolysis.

## Conclusions

Until far, research on non-cellular processes in soils has focused on CO_2_ emissions from sterilised soils (e.g. **9, 10, 11, 14, 15**). We open a “black box” by providing the first evidence that citrate and glucose in sterile soils trigger non-cellular convergent reactions that lead to the formation of several Krebs cycle intermediates, as well as additional fermentation products such as acetate and formate. These observations extend those of Bouquet et al. **(14)** who demonstrated a series of time-ordered reactions controlled by the nature of the substrate in sterile soils. Together, these results challenge the dogma of the cell as the minimal unit for carrying out metabolic reactions by showing that, under mild conditions (20°C, pH 6.2) in soils, a network of non-cellular reactions recapitulates most of the intermediates of the Krebs cycle.

Strengthening the idea of a real chemical continuity between prebiotic chemistry and biochemistry **(8)**, our observations bring up a number of issues not only regarding the diversity of these non-cellular metabolic networks (e.g. respiration, glycolysis, fermentation) but also their fundamental architecture (e.g. connection between metabolic intermediates, possible shunting of pathways). This will be pursued using specific positional labelling to track isotopomers and isotopologues as well as fluxomic approaches to study the kinetics and potential cyclisation of these processes.

From a biogeochemical perspective, while previous estimates suggest that EXOMETs could contribute significantly to CO_2_ emissions in contemporary soils **(10, 15)**, our study indicates that they could also contribute to CO_2_ emissions in anaerobic environments such as marine or freshwater sediments, through non-cellular fermentation.

## Materials and Methods

Supplementary methods appear in *SI Appendix*.

## Supporting information

Supporting information for Long lasting non-cellular reactions in sterile soils recapitulate most of the intermediates of the Krebs cycle

## Data Availability

GC-MS and GC-C-IRMS data are available in Figshare: https://figshare.com/s/5470d388bbc43f84ff69

## Acknowledgments

Clémentin Bouquet was supported by PhD fellowship from the French Ministry of Education and Research. The authors thank the financial support of (i) the CNRS through the Mission pour les Initiatives transverses et interdisciplinaires (MITI) and through Projet Exploratoire Premier Soutien (PEPS) and the funding of the ISO-EXOMET and EXCEED projects, (ii) CAP 20-25 Isite of the University Clermont Auvergne and the funding of the EXOMET project. The author(s) would like to thank SILVATECH (Silvatech, INRAE, 2018. Structural and functional analysis of tree and wood Facility, doi: 10.15454/1.5572400113627854E12) from UMR 1434 SILVA, 1136 IAM, 1138 BEF and 4370 EA LERMAB from the research center INRAE Grand-Est Nancy for it contribution to isotopic analysis and methodological development. SILVATECH facility is supported by the French National Research Agency through the Laboratory of Excellence ARBRE (ANR-11-LABX-0002-01)

