## Supporting information for Long lasting non-cellular reactions in sterile soils recapitulate most of the intermediates of the Krebs cycle for "Long lasting non-cellular reactions in sterile soils recapitulate most of the intermediates of the Krebs cycle"

**This file includes:**

Supporting Text
SI references

### Supporting Information Text

#### Methods

*Soil microcosm preparation procedure*

Soil characteristics, sampling, sterilisation and experimental design are described in detail in Bouquet *et al.* (2025). Briefly, non-sterilised soil (LS) and irradiated soil (IS) microcosms consisted of 10 g sieved soil samples (dry mass) were placed in 120 mL sterile glass vials capped with butyl rubber stoppers and sealed with aluminum crimps. Sterile solutions of uniformly labelled (*i.e.* all labelled positions) [$^{13}C_6$]-glucose (CAS: 110187-42-3, Cambridge Isotope Laboratories) and [$^{13}C_6$]-citrate (CAS: 287389-42-9, Sigma-Aldrich) or unlabelled glucose (CAS: 50-99-7, Sigma-Aldrich) and unlabelled citrate (CAS: 77-92-9, Sigma-Aldrich) were prepared. Five different substrate treatments, each with four replicates, were applied to sterilised soils (pH = 6.2): no substrate addition (IS), or addition of [$^{13}C_6$]-labelled citrate ($^{13}$C-IS), unlabelled citrate ($^{12}$C-IS), [$^{13}C_6$]-labelled glucose ($^{13}$G-IS), or unlabelled glucose ($^{12}$G-IS). Labelled substrates (0.05 % atoms) and unlabelled substrates were amended for a final concentration of glucose and citrate in a soil microcosm of 3 mgC. $g^{-1}$. All microcosms were flushed with sterilised free $CO_2$ gas (80 % $N_2$, 20 % $O_2$) moistened with sterilised water (for (LS) and (IS) microcosms), sterilised glucose or citrate solution (for ($^{12}$G-IS) and ($^{12}$C-IS) microcosms) or sterilised $^{13}C_6$-glucose and $^{13}C_6$-citrate solutions (for ($^{13}$G-IS) and ($^{13}$C-IS) microcosms) to a water content of 30 %. All microcosms were incubated in the dark at 20 °C for 163 days. Vials were sampled at various intervals (t= 0.2, 3, 6, 17, 23, 66, 100, 163 days) during the incubation. All manipulations were carried out under sterile conditions and each sampling was destructive to avoid contamination of the soil microcosms. The sterility of sterilised soils was confirmed (Bouquet *et al.* 2025).

*Exometabolite extraction from soils*

The water-extractable fraction of exometabolites < 3 kDa (WEM) of soil samples was obtained by Bouquet *et al.* (2025) using the method developed by Jenkins *et al.* 2017 with minor modifications. Briefly, four grams of soil were extracted under sterile conditions in 24 mL of sterile MilliQ water at 4°C for 1 h with shaking on an orbital shaker (Stuart SB3). The tubes were then centrifuged at 4 °C for 5 min at 3,200 × g. The supernatants were collected in fresh 50 mL conical tubes and centrifuged again. The supernatants were then filtered using Swinnex filtration units fitted with 25mm diameter GF/F (Whatman) previously grilled at 450°C for 4h and rinsed with 30 mL of sterile milliQ water. These water extractable metabolites were frozen at -20 °C, lyophilised to dryness (Christ ALPHA 1-2 LDplus) and resuspended in 1.5 mL ultrapure water (MilliQ system). The samples were filtered on a centrifugal filter (Amicon ultra-4, 3 kDa) previously rinsed using 4 mL sterile 0.1N NaOH and 4 mL sterile MilliQ water. After centrifugation for 45 min (7,500 g; 4°C), the WEM were lyophilised again and stored at -80°C until analysis

*Analyses of WEM using ion chromatrography coupled to mass spectrometry (IC-MS)*

Sample preparation: Each tube of dry sample was thawed for two hours at room temperature before being filled in 1 mL Eppendorf® tube containing ultrapure water (PureLab Veolia Water®). To obtain a processable signal, samples were diluted 100-fold in ultrapure water (PureLab Veolia Water®).

Technical settings: Standards for carboxylic acid compounds were purchased from Sigma-Aldrich, analytical grade standards quality (Cis-aconitic CAS 585-84-2; Citric CAS 77-92-9; Succinic CAS 110-15-6; Fumaric CAS 110-17-8; Malic CAS 6915-15-7; Oxaloacetic CAS 328-42-7; Alpha-ketoglutaric CAS 328-50-7; Formic CAS 64-18-6; Acetic CAS 64-19-7. The eluent (KOH) and sample solutions were prepared using ultrapure water from a (PureLab Veolia Water®).
Retention times and detection/quantification limits were determined as follows:

| | Mass (in m/z) | Retention time (in min) | Limit of detection (in ppm) | Limit of quantification (in ppm) | Limit of detection (in µg.g$^{-1}$ of dry soil) | Limit of quantification (in µg.g$^{-1}$ of dry soil) |
|---|---|---|---|---|---|---|
| Oxaloacetic acid | 131 | 24.8 | 1.0 ± 0.2 | 4.0 ± 0.2 | 0.375 ± 0.075 | 1.5 ± 0.075 |
| Alpha-ketoglutaric acid | 145 | 25.0 | 0.3 ± 0.2 | 1.0 ± 0.2 | 0.1125 ± 0.075 | 0.375 ± 0.075 |
| Succinic acid | 117 | 26.2 | 0.2 ± 0.1 | 0.5 ± 0.1 | 0.075 ± 0.0375 | 0.1875 ± 0.0375 |
| Malic acid | 133 | 26.2 | 0.05 ± 0.01 | 0.10 ± 0.01 | 0.01875 ± 0.00375 | 0.0375 ± 0.00375 |
| Fumaric acid | 115 | 28.3 | 0.05 ± 0.01 | 0.10 ± 0.01 | 0.01875 ± 0.00375 | 0.0375 ± 0.00375 |
| Cis-aconitic acid | 173 | 32.8 | 0.05 ± 0.01 | 0.10 ± 0.01 | 0.01875 ± 0.00375 | 0.0375 ± 0.00375 |
| Citric acid | 191 | 32.0 | 0.05 ± 0.01 | 0.10 ± 0.01 | 0.01875 ± 0.00375 | 0.0375 ± 0.00375 |
| Formic acid | 45 | 10.0 | 5 ± 1 | 16 ± 1 | 1.875 ± 0.375 | 6 ± 0.375 |
| Acetic acid | 59 | 8.4 | 3.1 ± 0.9 | 9.1 ± 0.9 | 1.1625 ± 0.3375 | 3.4125 ± 0.3375 |

The limit of detection (LD) was estimated with a signal to-noise ratio of 3:1, while the limit of quantification (LQ) was obtained using a signal-to-noise ratio of 9:1 (Shrivastava and Gupta 2011). Calibration curves were also generated for compounds to validate the limits of detection and limits of quantification obtained through the signal-to-noise method. Detection and quantification limits in ppm correspond to limits measured in solution on the instrument. Their correspondence in µg.g$^{-1}$ of dry soil in the soil studied is then calculated by multiplying the value in ppm by 0.375, corresponding to the ratio between the extraction volume (1.5mL) and the quantity of soil extracted (4g) (see table above). IC-MS analyses were carried out on a Thermo-Fisher Scientific ICS-6000 interfaced to a simple quadrupole mass spectrometer (ISQ-EC - Thermo Scientific) using SIM mode after Scan mode for pre-screening. The pre-column was a CG-11-HC and the column was a Dionex IonPac AS-11-HC-4µm 2×250 mm column, with the KOH step gradient starting at 1 mM KOH from 0 to 5 min, 10 mM KOH from 5.1 to 20 min, 30 mM KOH from 20.1 to 25 min, 60 mM KOH from 25.1 to 31 min. The latter concentration was held until 35 min, and the column was re-equilibrated at 1 mM KOH at 35.1 min at a flow rate of 0.36 mL min-1 and temperature of 40°C. Instrument control, data acquisition and processing were performed via Chromeleon 7 Software, version 7.2.10.

*Analyses of WEM using gas chromatography (GC) coupled to mass spectrometry (MS) or combustion isotopic ratio mass spectrometry (C-IRMS)*

Chemical derivatisation of samples: Prior to GC analyses, the WEM samples were chemically derivatised (Wang *et al.* 2008) into volatile, low-polarity, and thermally stable molecules because Krebs cycle intermediates contain carboxyl and hydroxyl groups, which are less volatile and highly polar (Li *et al.* 2023). Each dry sample of sterilised soil amended with unlabelled substrate ($^{12}$G-IS and $^{12}$C-IS) or labelled substrate samples ($^{13}$G-IS and $^{13}$C-IS) was filled into 200 µL of a mixture of methanol (Honeywell Riedel de Hoën LC-MS Chromasolv TM >=99.9%) and 10 % sulfuric acid (Honeywell Fluka ACS reagent 95-97%) and collected in reaction tube (Wheaton 1mL Grad V-Vial®) before being vortexed 5 s and placed in a dry bath (Stuart block heater SBH130DC) at 60°C for 1 h. After cooling at room temperature, 200 µl (for GC-C-IRMS) or 300 µL (for GC-MS) hexane was added and samples were vortexed 10 s before centrifugation at 1000 rpm for 3 min (Eppendorf centrifuge 5702). The supernatant (200 µL) was collected and transferred to 1.5 mL vials (11 x 32 mm, Fisher scientific) with 250 µL inserts (Agilent glass with polymer feat and screw cap Fisherbrand 8 mm black UC Sil/PTFE 1.3mm). After GC-MS analyses, caps were changed and samples were frozen at -20°C for subsequent analyses in GC-C-IRMS analyses.

Technical settings: GC-MS and GC-C-IRMS analyses were performed on an Agilent® DB-35ms column, model number 122-3832E (30 m, 0.250 mm, 0.25 µm film thickness). Metabolite retention times were determined using GC-MS, on a GC 6890N (Agilent®) equipped with an MSD 5973N mass detector (Agilent®). An electron ionization system with an ionization energy of 70 eV was used for GC-MS detection, and helium gas was used as carrier gas at a constant flow rate of 1.1 mL/min. Hexane was used as the wash solvent. An injection volume of 1 µL of sample was considered in the analysis. The initial temperature of the front inlet was 260 °C and the split ratio 10:1. The injector and MS transfer line temperature was set at 280 °C. The oven temperature was programmed from an initial temperature of 50 °C (hold for 1 min) to a final temperature of 320 °C at an increasing rate of 6 °C/min to 150 °C, 20 °C/min to 250 °C and 30 °C/min to 320 °C (hold for 5 min). The acquisition was performed in scan mode, with a solvent delay of 2.5 min. Metabolites with mass spectrum fragments between 35 and 350 (m/z) were analysed. Non-volatile Acid Standard Mix (analytical grade, Sigma-Aldrich) was used to determine the retention time of the compounds. For cis-aconitic the NIST23 (2023) library was used (Sisco and Moorthy 2020), and for malic and alpha-ketoglutaric pure standards (CAS 636-61-3; CAS 328-50-7 Sigma-Aldrich, analytical grade) were used. Data from GC-MS were analysed using software Agilent Mass Hunter Workstation Unknown Analysis version 12.1.
The relative abundance of carbon isotopes ($^{13}C/^{12}C$) of targeted metabolites were determined using GC-C-IRMS for $^{12}$G-IS, $^{12}$C-IS, $^{13}$C-IS and $^{13}$G-IS samples using a gas chromatograph (Agilent 7890A) coupled to a continuous-flow isotope ratio mass spectrometer (Elementar UK, Cheadle, UK) via a

combustion module. This setup is available at SILVATECH (Silvatech, INRAE, 2018. Structural and functional analysis of tree and wood facility, doi: 10.15454/1.5572400113627854E12). A 10 µL syringe was used to inject the sample into the injector in splitless mode at 250 °C, and the final injection volume was 2 µL. After separation, the compounds were combusted at 850°C in a quartz tube filled with copper oxide. The resulting gases were then analysed using an isotope ratio mass spectrometer (Isoprime).

We also determined the $^{13}C/^{12}C$ ratio for unlabelled glucose and citrate dry powders (both substrates showed a close initial isotopic signature without labelling (-10.68 ±0.2 ‰ for citrate, and -11.64 ± 0.2 ‰ for glucose) by placing tin capsules containing dry material in an elemental analyzer (vario ISOTOPE cube, Elementar, Langenselbold, Germany) coupled, via a gas box interface, to the continuous flow isotope ratio mass spectrometer (Isoprime100, IRMS, Elementar UK, Cheadle, United Kingdom) The sample was burned at 1025°C in excess of oxygen. Then, the NOx was reduced using a quartz tube filled with copper heated at 650°C. $CO_2$ was trapped at 35°C by an adsorption column while $N_2$ passed through the thermal conductivity detector (TCD). Next, $CO_2$ was released from the adsorption column at 225°C.

Elementary gases were analyzed and detected by isotope ratio mass spectrometry, Isoprime100 IRMS (Cheadle, United Kingdom). The $\delta^{13}C$ values were expressed as delta values in ‰ relative to the isotope ratio of the Vienna Pee Dee Belemnite (VPDB) standard. The d13C uncertainty of measurements is 0.3 ‰ (2σ) (i.e. not pure molecule).

Substrates showed an identical isotopic signature after labelling (Isotopic abundance of $^{13}C$=0.05, i.e. theoretical delta $^{13}C$ = 3707.7 ‰, U-$^{13}C_6$-Glucose or U-$^{13}C_6$-Citrate). Note that malic acid was not detected by GC-C-IRMS because the derivatisation procedure destroyed this chemical, resulting in the formation of succinic acid (26.6 ± 0.7 %) and, to a lesser extent, fumaric acid (10.8 ±0.4 %).

GC-C-IRMS were analysed using softwares Isoprime100 IRMS (Cheadle, United Kingdom) and IonVantage version 1.7.3.0. The results were expressed as delta values (‰) relative to VPDB.

s

*Statistical analyses:*

All statistical analysis and tests were performed using GraphPad Prism version 8.0.1 (GraphPad Software, Inc). A paired t-test two tailed p. value (n=4) was used firstly in order to compare the insignificantly impact of labelling in concentration of metabolites. For all analysed metabolites, labelling did not affect significantly metabolite concentration. A paired t-test two tailed p. value (n=8) was then applied to compare the concentration of metabolites according to whether or not citrate or glucose was added to the soil. Only for cis-aconitic, as the concentration values in the unamended soil were mostly equal to or below the quantification threshold, we used a one-sample t-test against a theoretical mean corresponding to the quantification threshold. Finally, a paired t-test (n=4) two tailed p. value was also applied to compare isotopic ratios ($\delta^{13}C$)) between samples amended with labelled or unlabelled substrates.

*Data availability:*

GC-MS and GC-C-IRMS data are available in Figshare: https://figshare.com/s/5470d388bbc43f84ff69
